# Muscle stem cell function is impaired in absence of Talpid3 - a gene required for primary cilia formation

**DOI:** 10.1101/2022.10.14.512102

**Authors:** Victor Martinez-Heredia, Danielle Blackwell, Sujith Sebastian, Timothy Pearson, Gi Fay Mok, Laura Mincarelli, Charlotte Utting, Leighton Folkes, Ernst Poeschl, Iain Macaulay, Ulrike Mayer, Andrea Münsterberg

## Abstract

Skeletal muscle stem cells (MuSC) are crucial for tissue homeostasis and repair after injury. Following activation, they proliferate to generate differentiating myoblasts. A proportion of cells self-renew, re-enter the MuSC niche under the basal lamina outside the myofiber and become quiescent. Quiescent MuSC have a primary cilium, which is disassembled upon cell cycle entry. *Ex vivo* experiments suggest cilia are important for MuSC self-renewal, however, their role in muscle regeneration *in vivo* remains poorly understood. Talpid3 (TA^3^) is essential for primary cilia formation and Hedgehog (Hh) signalling. Here we use tamoxifen-inducible conditional deletion of TA^3^ in MuSC (iSC-KO) and show that regeneration is impaired in response to cytotoxic injury. Repeat injury exacerbates the regeneration phenotype in TA^3iSC-KO^ mice, indicating depletion of MuSCs. Single cell transcriptomics of MuSC progeny isolated from myofibers identifies components of several signalling pathways, which are deregulated in absence of TA^3^, including Hh and Wnt. Pharmacological activation of Wnt restores muscle regeneration, while purmorphamine, an activator of the Smoothened (Smo) co-receptor in the Hh pathway, has no effect. Together, our data suggest that TA^3^ and primary cilia are important for MuSC self-renewal, and that pharmacological treatment can efficiently restore muscle regeneration.

## Introduction

Primary cilia are non-motile microtubule-based structures found on many cells. They serve as a cellular antenna for multiple signalling pathways and are implicated in numerous developmental disorders. The importance of primary cilia for Sonic hedgehog (Hh) signalling in vertebrates was first uncovered in a mouse mutagenesis screen, which identified genes encoding intraflagellar transport (IFT) proteins ^1^ required to build and maintain primary cilia. The sequestration of Hh pathway components into the cilium regulates the down-stream processing of GLI effector proteins from transcriptional repressors - in absence of the ligand - into activators when the pathway is active ^(2–4^. Smoothened (Smo), a seven-pass transmembrane protein and Hh co-receptor, also moves into the ciliary membrane depending on pathway activity. A conserved ciliary localisation motif has been identified ^5,6^.

Many ciliary proteins have now been characterized and mutations in some of these are associated with diseases in human, collectively termed ciliopathies ^7^. Talpid^3^ (TA^3^) is a critical component of primary cilia. The protein is located to the distal end of centrioles and necessary for docking of the basal body with the apical cell membrane and thus, for primary cilia formation ^8 9^. The TA^3^ gene was first identified in chicken and shown to be essential for embryonic patterning and myogenesis ^10^. The defects seen in the original TA^3^ chicken mutant ^11^ include features that can be attributed to abnormal Hh signalling, such as limb and neural tube defects. These were also phenocopied in a mouse knockout, where exons essential for TA^3^ protein function were deleted ^8, 12^.

The Hh pathway is the most investigated cilia-related signalling pathway, however, there is extensive crosstalk between Wnt and Hh signalling pathways, which share common regulators - including GSK3ß, CK1αand Smoothened (Smo). Both pathways are linked to primary ciliogenesis and depending on the context, they act cooperatively or antagonistically ^13 14 15 16^

Quiescent muscle stem cells (MuSC) have a primary cilium, which is disassembled upon activation and cell cycle entry and reassembled in self-renewing MuSCs ^17^. Knock-down of IFT88 in C2C12 myoblasts leads to ablation of ciliogenesis and reduced expression of p27, a marker for quiescence ^18^. In mice, conditional deletion of IFT88 in MuSCs, impairs recovery of overall muscle strength and alters the expression of cell-cycle-related genes ^19^. Repression of Hh signalling seems to be important in maintaining MuSCs quiescence in Go ^19, 20^. This requires the repressor form of GLI3, since conditional deletion of GLI3, or inhibition of its processing into GLI3R, leads to cell cycle entry ^21^. Together this suggests that Hh signalling regulates MuSCs activation and proliferation. However, the role of primary cilia as a nexus for signalling pathways to modulate skeletal muscle regeneration *in vivo* in response to injury has not yet been investigated.

Here we use the floxed allele of TA^3 8, 12^, to generate an inducible conditional knock-out in MuSCs, termed TA^3iSC-ko^, using PAX7-Cre^ERT2^. In adult mice Pax7, and thus Cre^ERT2^, is specifically expressed in MuSCs and triggers recombination of the floxed TA^3^ locus in these cells after tamoxifen administration ^22, 23^. Using cardiotoxin (ctx)-induced injury, we show that TA^3^ function is required for efficient muscle repair. In TA^3iSC-KO^ mutants, fiber diameters were smaller and repeat injury enhanced the phenotype, suggesting MuSC self-renewal is affected

We used two reporter alleles, Pax7-ZsGreen ^24^ or Rosa-fl-stop-fl-Td-Tomato, to FACS isolate MuSCs and their progeny either *in vivo* or from *ex vivo* cultured myofibers, followed by transcriptomics analysis. This revealed effects on ECM related genes and de-regulated signalling pathways in absence of TA^3^; in particular, many Wnt ligands were downregulated. Consistent with the latter, we find that muscle regeneration was rescued by co-injection of BIO (6-bromoindirubin-3’- oxime), an activator of ß-catenin-dependent Wnt signalling, but not by purmorphamine (pur) ^25^, an activator of the Hh co-receptor, Smo. We propose that modulation of signalling via the primary cilium promotes MuSC self-renewal, and that this might be important in muscle disease or in ageing when MuSC function declines.

## Results

### TA^3^ is necessary for primary cilia formation in MuSCs

To examine the requirement of Talpid3 (TA^3^) for primary cilia formation in MuSCs, we used a conditional allele of the TA^3^ gene ^12^ with the Pax7^CreERT2^ driver ^22, 23, 26^. Homozygous TA^3flox/flox^/ Pax7^CreERT2^ animals (TA^3iSC-ko^) lack expression of the gene in MuSCs. Heterozygous TA^3flox/+^ litter mates or animals lacking Pax7^CreERT2^ were used as controls.

Isolated myofibers were cultured for 0, 48 and 96 hours and immunostained using Arl13b and Pax7 antibodies to detect primary cilia in MuSC (Fig. 1A). At 0 and 48 hours the percentage of Pax7 positive cells with cilia was similar in TA^3SC-KO^ and control fibers. This was expected, as primary cilia are only disassembled upon entry into the cell cycle ^17^, which occurs >40 hours after activation. After 96 hours in culture, MuSCs have undergone multiple rounds of cell division, a proportion of cells reexpress Pax7, re-enter the stem cell niche and re-assemble a cilium. At this point 60% of Pax7 expressing cells had a primary cilium in control myofibers, similar to previous reports ^17, 21^. In TA^3iSC-KO^ mutants there were fewer Pax7 positive (Pax7^+ve^) cells with a primary cilium although the number of Pax7^+ve^ cells does not significantly change (Fig. 1B, C).

**Figure 1:**
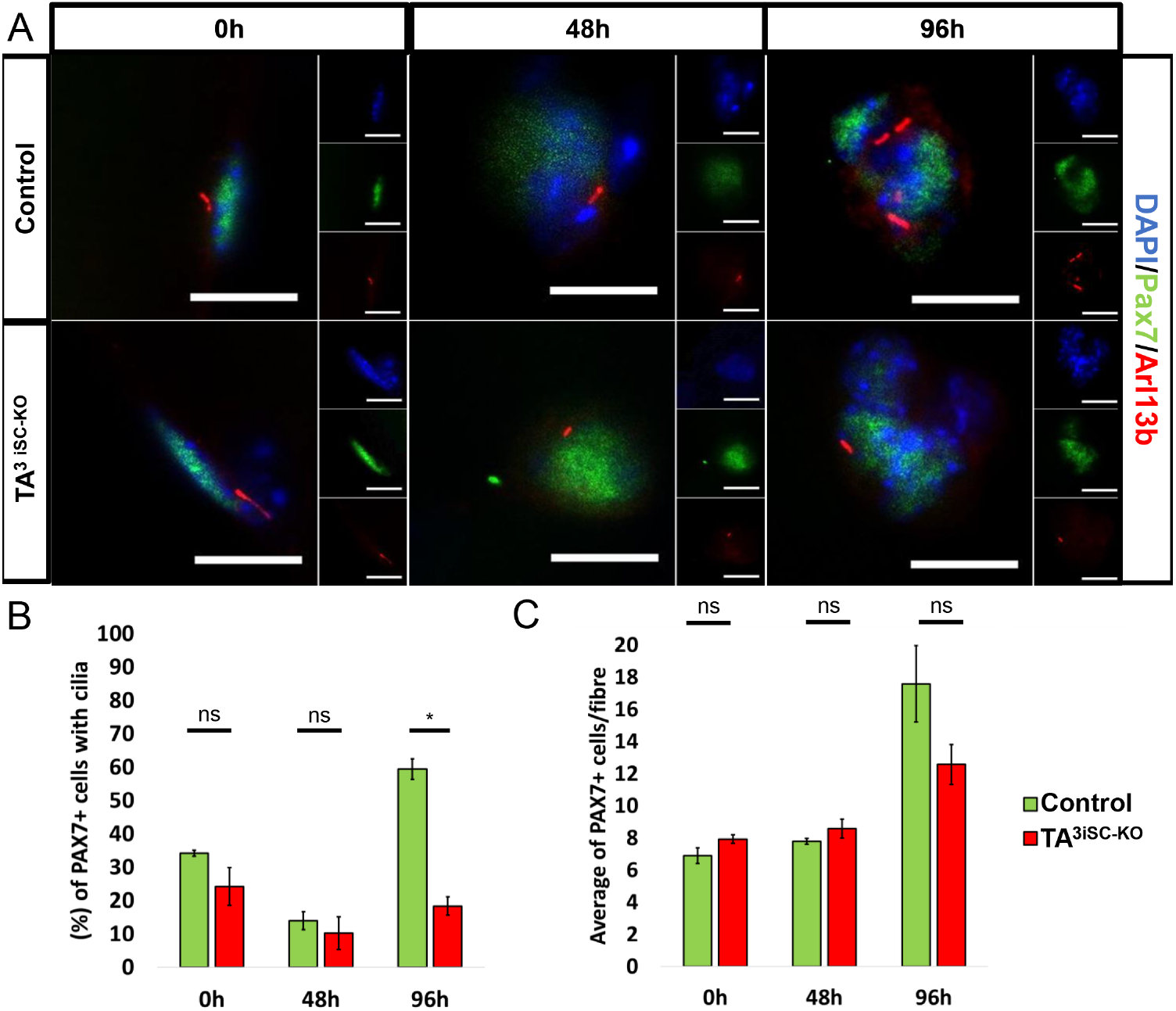
Cilia assembly in MuSCs requires TA^3^. (A) Representative images of EDL muscle fibers from TA^3iSC-ko^ and control mice at 0h, 48h and 96h. Immunofluorescence detects Pax7 expressing MuSCs (green) and primary cilia using Arl13b (red), DAPI (blue) stains nuclei (size bars = 10 μm). (B) The percentage of Pax7 cells positive for Arl13b in fibers from TA^3iSC-ko^ and control mice, *p < 0.05, n=3 mice and >10 fibers per mouse. (C) The average number of Pax7 positive cells per fiber after 0h, 48h and 96 h in culture, control (green) and TA^3iSC-ko^ (red) is not significantly different.

### MuSCs lacking TA^3^ have a proliferation defect

To determine cellular defects arising from lack of TA^3^, control and TA^3iSC-KO^ myofibers were cultured *ex vivo* to characterise activation, proliferation and differentiation of MuSCs in their physiological niche (Fig. 2A). Myofibers were analysed at different time points by immunostaining for Pax7, Myf5, MyoD and MyoG, and nuclei were counted using DAPI. Pax7 expression marks quiescent MuSCs. Once cells are activated, they express Myf5, followed by MyoD and MyoG, which indicates differentiation. A proportion of cells maintains or re-expresses Pax7 and downregulates MyoD to self-renew the stem cell pool ^27^. At 0, 24 and 48 hours we detected no differences in either the total number of cells or the proportion of cells expressing different markers (Supplementary Fig. 1, Supplementary Fig. 2). However, from 72 hours TA^3iSC-KO^ mutant myofibers had fewer MuSC-derived cells compared to controls. The number of cell clusters per myofiber was not affected, but there were fewer cells per cluster (Fig. 2B), suggesting a defect in MuSC activation or proliferation. Caspase 3 staining showed no difference between TA^3iSC-KO^ and control myofibers suggesting the reduced cell number was not due to an increase in apoptosis (data not shown). In TA^3iSC-KO^ mutants, the proportion of cells positive for Pax7, or MyoD, or both was not affected (Fig. 2C, C’). This is also the case at 96 hrs (Supplementary Fig. 1, Supplementary Fig. 2). Immunostaining for MyoD and MyoG showed no difference in the proportion of single and double positive cells (Fig. 2D, D’), indicating that myogenic differentiation was not impaired, although there were fewer cells overall.

**Figure 2:**
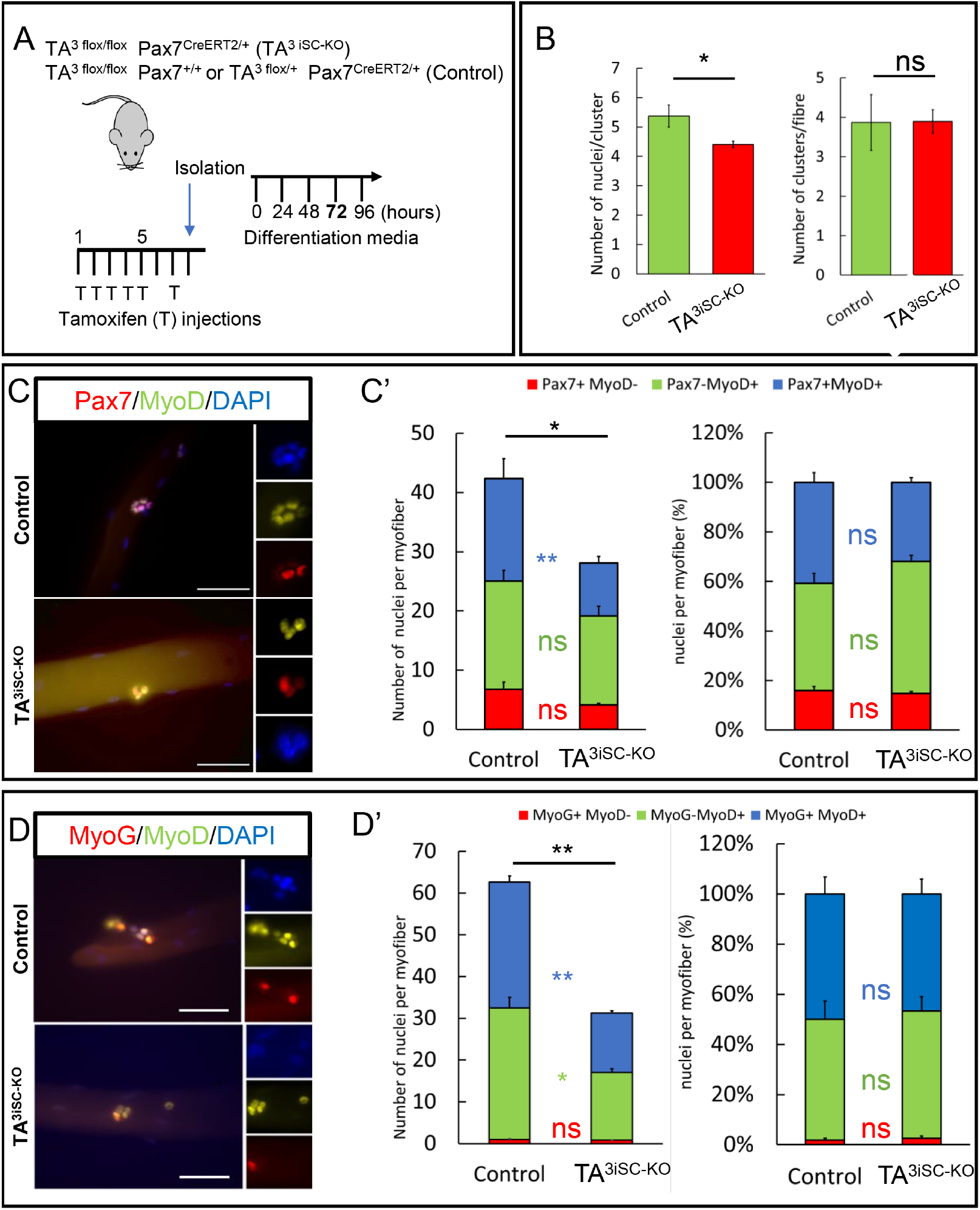
Efficient proliferation of MuSCs requires TA^3^. (A) Schematic illustration of tamoxifen injection regime followed by *ex vivo* culture of myofibers for 72 hours. Genotypes of mutant and control mice are shown. (B) Clusters of MuSCs were counted and plotted as the total number of nuclei per fiber or the number of clusters per fiber. Overall fewer MuSC-derived cells were present inTA^3iSC-KO^ compared to controls. (C) Pax7 and MyoD or (D) MyoD and MyoG immunostaining of MuSC progeny on myofibers isolated from control or TA^3iSC-KO^ mice (size bar = 50 μm). (C’) Graphs show the total number of nuclei per myofiber in control and TA^3iSC-KO^ fibers, or the percentage of Pax7^+^ (red), MyoD^+^ (green) or Pax7^+^/MyoD^+^ (blue) double positive cells *p < 0.05 **p<0.01, n=9. (D’) Graphs show the total number of nuclei per myofiber in control and TA^3iSC-KO^ fibers, or the percentage of MyoG^+^ (red), MyoD^+^ (green) or MyoG^+^/MyoD^+^ (blue) double positive cells *p < 0.05 **p<0.01, n=9.

### Loss of TA^3^ in MuSC impairs skeletal muscle repair after injury

Next, we determined the effect of primary cilia loss on the ability of skeletal muscle to repair *in vivo*,following injury induced by cardiotoxin (ctx) injection. Deletion of TA^3^ was induced in MuSCs prior to injury and hindlimb muscles were dissected on day 5, 10 or 15 post-injury. Histology of transverse cryosections and nidogen staining of basement membranes showed disorganised myofibers at day 10 post-injury (Fig. 3A, B). DAPI staining confirmed centrally located myonuclei in regenerated myofibers compared to non-injured control muscles, where myonuclei are found in the periphery (Fig. 3B). In uninjured muscles, the distribution of Feret diameters was identical in control and TA^3iSC-KO^ myofibers (Fig. 3C, D). However, following injury regenerating TA^3SC-KO^ myofibers had smaller average Feret diameters compared to controls at 10 (Fig. 3C, D). This shows muscle regeneration is impaired in absence of TA^3^.

**Fig. 3:**
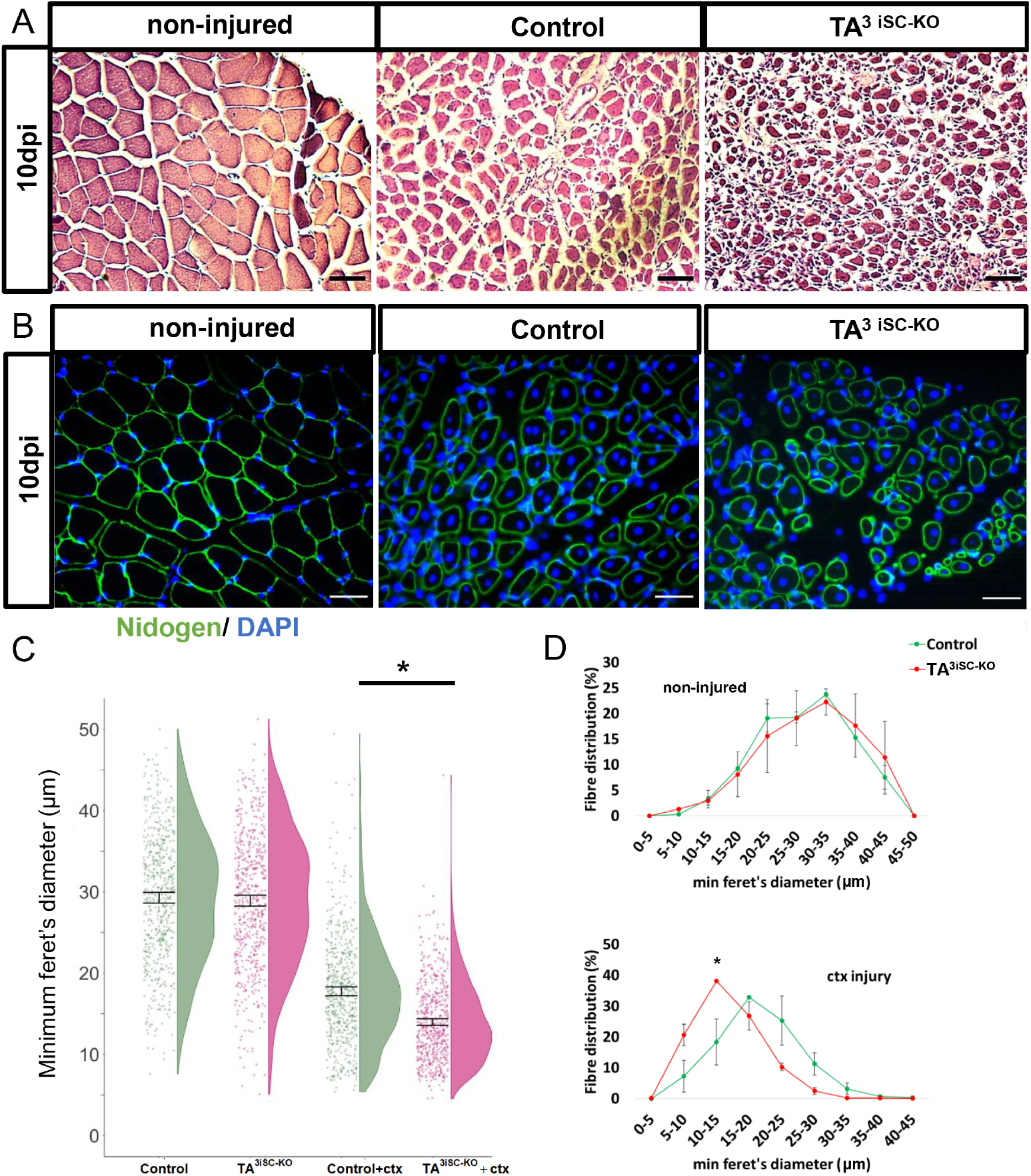
Regeneration of myofibers is impaired after deletion of TA^3^ in MuSCs. (A) Histology images of *tibialis anterior* muscle transverse sections stained with Hematoxylin and Eosin. (B) Representative images of TA transverse sections, nidogen staining visualises basement membrane (green), DAPI stains nuclei (blue). Sections from non-injured, and injured control and TA^3iSC-KO^ muscles are shown as indicated, 10dpi = 10 days post-injury (size bar = 50μm). (C) Raincloud plots showing size distribution of myofiber minimal Feret diameters in control (green) and TA^3iSC-KO^ (red) mice, non-injured or after cardiotoxin (ctx) injury, *p < 0.05. (D) Histograms of minimal Feret diameters of myofibers in control (green) and TA^3iSC-KO^ (red) *tibialis anterior* muscles with (bottom) or without (top) injury, n=3 mice per genotype, *p < 0.05.

### Regeneration is more severely compromised in TA^3iSC-KO^ after repeat injury

To determine whether self-renewal of MuSCs is affected we used a repeat injury, where a second injury was applied after a 25-day recovery period (Fig. 4A). Histology was similar in control muscles 10 days after repeat injury compared to single injury. However, TA^3iSC-KO^ mutants showed more severe fibrosis after repeat injury compared to single injury (Fig. 4B). Furthermore, myofiber diameters were smaller in TA^3iSC-KO^ mutant compared to control muscles 10 days post repeat injury (Fig. 4C-E), and myofiber diameters were more reduced after repeat injury compared to single injury (5-10μm compared to 10-15μm) (Fig. 3, Fig. 4). This suggests that MuSC are depleted after a first injury, consistent with the notion that self-renewal is affected. In non-injured control and mutant muscles fiber diameters were similar.

**Fig. 4:**
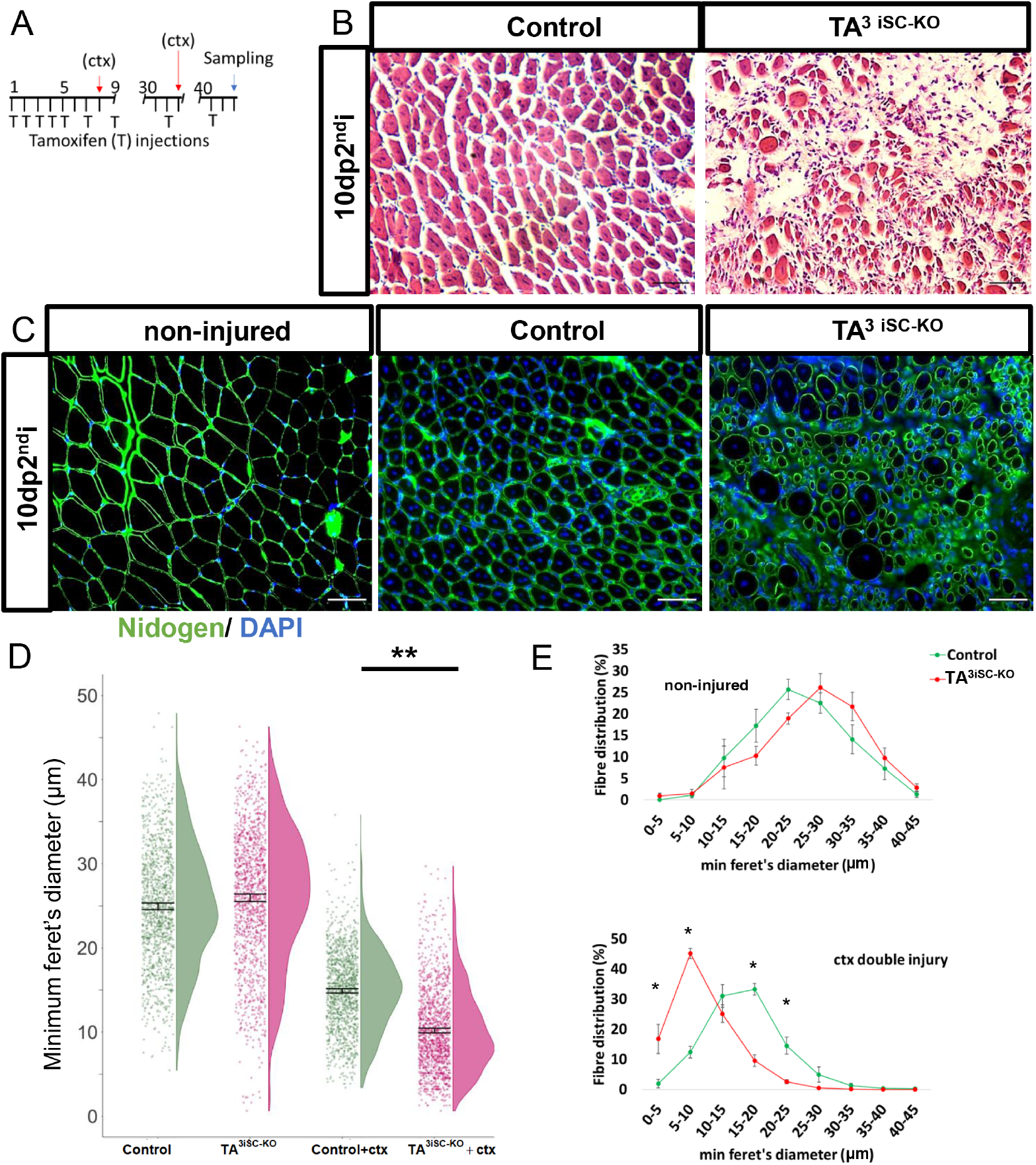
A second injury enhances regeneration defects in TA^3iSC-KO^ mice. (A) Schematic of the tamoxifen (T) and ctx injection regime. The first injury on day eight was followed by a second injection 25 days later. Analysis was performed ten days post-second injury (10dp2^nd^i). (B) Histology shows good recovery in controls but poor recovery in TA^3iSC-KO^ mice. (C) Transverse sections stained with nidogen (green) and DAPI (blue) at 10dp2^nd^i. Sections from non-injured and injured muscles from control and TA^3iSC-KO^ mice are shown as indicated (size bar = 50μm). (D) Raincloud plots showing the size distribution of myofiber minimal Feret diameters in control (green) and TA^3iSC-KO^ (red) mice, either non-injured or 10 days after the second injury (10dp2^nd^i) (ctx). (E) Histograms of minimal Feret diameters in control (green) and TA^3iSC-KO^ (red) *tibialis anterior* muscles without (top) or with repeat injury (bottom), n≥3 mice per genotype, *p < 0.05, **p < 0.01.

### Single cell analysis reveals deregulated signalling pathways in TA^3iSC-KO^ MuSCs

To determine the mechanisms underlying the loss of MuSCs after 72 hours in *ex vivo* cultures and the impaired muscle regeneration after injury, we established molecular profiles of MuSCs and their progeny. We first examined whether there are immediate effects on gene expression after TA^3^deletion and performed differential transcriptomics of quiescent MuSCs isolated from control and TA^3iSC-KO^mice. Pax7-Zs-green fluorescence ^24^ was used to FACS-isolate between 30000 and 50000 MuSCs from dissected hindlimb muscles, immediately following tamoxifen-induced TA^3^ deletion, for RNAseq. Principle component analysis did not clearly separate the control and TA^3iSC-KO^ samples and only few differentially expressed genes were identified, with 35 genes up-regulated and 27 genes down-regulated. GO-term analysis did not highlight specific pathways or processes that were enriched. Therefore, the loss of TA^3^ did not lead to significant molecular differences in quiescent MuSCs. We noticed, however, that several genes associated with ECM were differentially expressed, such as TIMP1, LAMB1 and ITGA8 (Supplementary Fig. 3).

To determine whether TA^3^-loss affects MuSC and their progeny as they go through the regenerative cycle, we performed single cell transcriptomics. Myofibers were cultured *ex vivo* (72 hrs) and Td-Tomato fluorescent MuSCs were isolated by FACS, sorted into plates and sequenced using Smart-Seq2. From three independent biological replicate samples each, for control and TA^3iSC-KO^ MuSCs, we retained 832 MuSCs after quality control. Downstream scRNA-seq analysis was performed using Seurat ^28^. The cells grouped into three clusters based on distinct transcriptional programs and expression of relevant marker genes (Fig. 5A, B). The top 40 genes identified as predominantly expressed in these three clusters are shown (Fig. 5B). A population of cells characterized by Pax7 was separated further into two distinct clusters (Fig. 5A, D). GO-terms related to cell proliferation were enriched in one of these Pax7 positive clusters and the top 10 differentially expressed genes included cyclin dependent kinase 1 (Cdk1) (Fig. 5C, D). This cluster was designated proliferating, pMuSC. The second cluster comprising Pax7 expressing cells was designated MuSC. Enriched GO-terms included extracellular matrix (ECM) organization and Fibronectin (Fn1) and Extracellular matrix protein 1 (Ecm1) were amongst the top 10 differentially expressed genes. The third cluster comprises differentiating cells characterized by MyoG expression. GO-terms were enriched for genes associated with myofibril assembly, striated muscle development and contraction. The top 10 differentially expressed genes include myogenin (MyoG), Actinin alpha3 (Actn3) and titin (Ttn). In both control and TA^3iSC-KO^ the different populations of MuSC progeny are distributed similarly across all clusters (Fig. 5A). This is consistent with our analysis of *ex vivo* cultured myofibers by immunostaining, where there was no change in proportions of cells labelled by Pax7, Myf5 or MyoG, although total cell number was reduced (Fig. 2, Supplementary Fig. 1, Supplementary Fig. 2).

**Figure 5:**
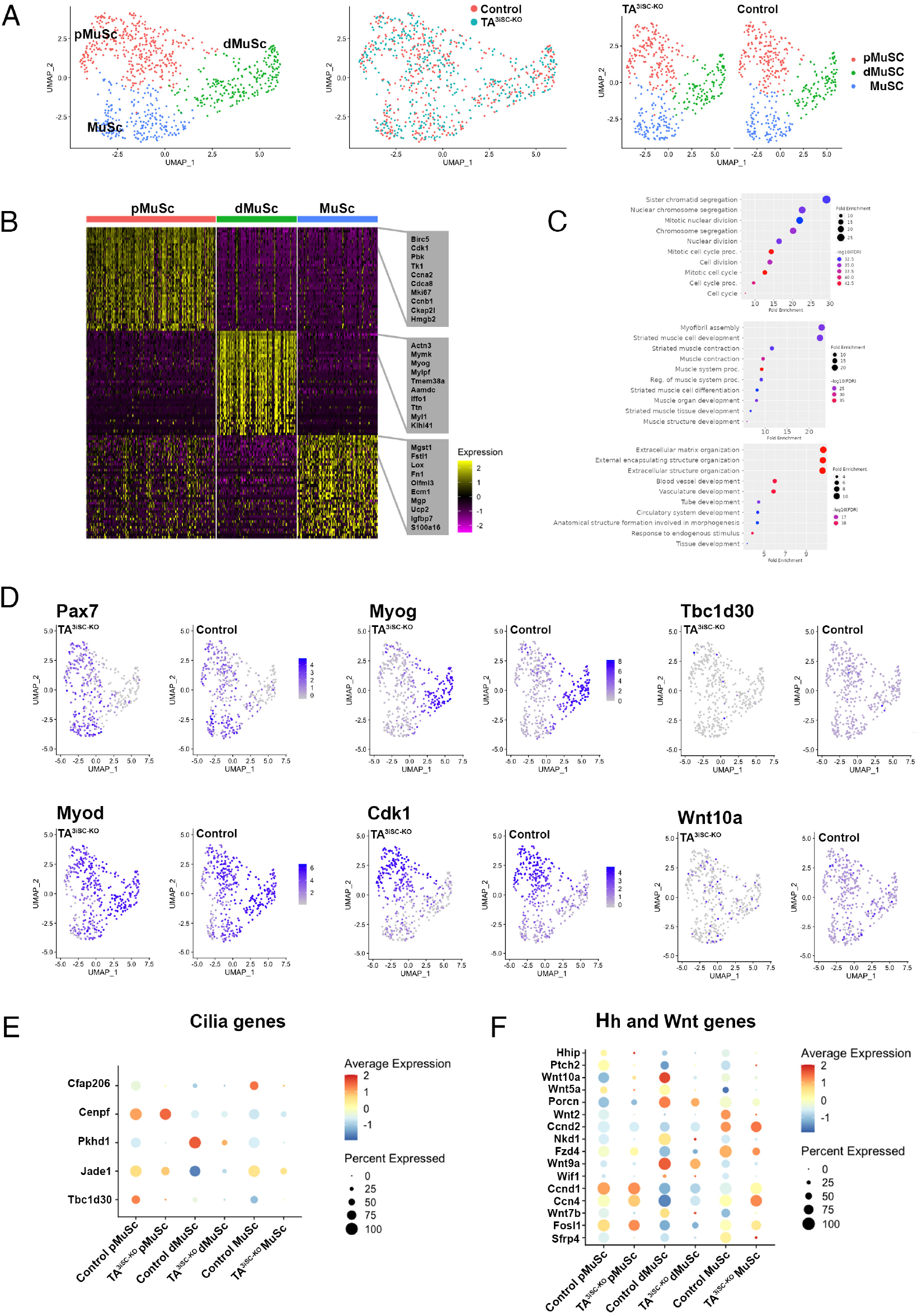
Loss of TA^3^ leads to deregulation of signalling pathways in MuSC progeny. (A) UMAP clustering of Td-Tomato MuSC progeny from *ex vivo* cultured myofibers identifies equal proportions of proliferating MuSC (pMuSC), differentiating MuSC progeny (dMuSC) and MuSC populations in control and TA^3iSC-KO^ myofibers. (B) Heatmap shows the top 40 most variably expressed genes between the three MuSC clusters, with the top 10 genes highlighted in grey. (C) Gene Ontology (GO) terms enriched in pMuSC, dMuSC and MuSCs populations are shown. (D) Expression of different marker genes across the three clusters in control and TA^3iSC-KO^ MuSC. Markers shown are Pax7, MyoD, MyoG, Cdk1, Tbc1d30 and Wnt10a. (E) Comparison of the expression of cilia genes and (F) Hedgehog (Hh) and Wnt signalling components across each of the three clusters.

The distribution of cells across each cluster was the same in control and TA^3iSC-KO^ populations (pMuSC: control = 188, TA^3iSC-KO^ = 183; dMuSC: control = 112; TA^3iSC-KO^ = 119; MuSC: control = 115; TA^3iSC-KO^ = 115). However, differential analysis revealed changes in gene expression between control and TA^3iSC-KO^. The expression of cilia associated genes was reduced in TA^3iSC-KO^ MuSCs. This includes Cilia and Flagella Associated Protein 206 (Cfap206) and TBC1 Domain family member 30 (Tbc1d30), a protein involved in regulation of cilia assembly (Fig. 5D, E). Examination of signalling pathways showed that genes involved in Hh signalling were reduced in MuSCs lacking TA^3^, including Hedgehog interacting protein (Hhip) and the Patched 2 receptor (Ptch2) (Fig. 5F). Furthermore, expression of many Wnt ligands was reduced in TA^3iSC-KO^ either across all three clusters (Wnt10a, Wnt5a, Wnt7b), or specifically in the Pax7 positive clusters, pMuSCs and qMuSCs (Wnt2, Wnt9a) (Fig. 5D, F). We also noticed that expression of both positive and negative regulators of Wnt signalling was reduced in TA^3iSC-KO^ MuSC progeny. Differentially expressed genes include porcupine (Porcn), which mediates Wnt palmitoylation required for activity, and naked cuticle homolog 1 (Nkd1) and secreted frizzled related protein 4 (Sfrp4), which inhibit Wnt signalling. Because components of Hh and Wnt signalling were deregulated in TA^3iSC-KO^ cells, we tested whether pharmacological activation of these pathways could rescue muscle repair after injury.

### Pharmacological inhibition of GSK3ß-kinase restores muscle regeneration in TA^3iSC-KO^ mice

The primary cilium is a nexus for Hh signalling and it has been shown that the co-receptor, Smoothened (Smo), which regulates downstream processing of Gli proteins to generate transcriptional activators, exhibits signalling competency in the absence of ciliary accumulation ^29^. Thus, we used purmorphamine (pur), a pharmacological activator of Smo, to determine whether this could restore the regenerative defect in TA^3iSC-KO^ mice. We found, however, that after co-injection of pur with ctx myofiber diameter remained reduced on day ten in injured TA^3iSC-KO^ muscle compared to controls (Fig. 6B-D). Expression of Hh responsive genes, Gli1 and Ptch1, was increased in biopsies of whole muscles after pur treatment (Fig. 6E). The same was observed in MuSCs isolated by FACS (Fig 6F, Supplementary Fig. 4). Thus, TA^3iSC-KO^ MuSCs can activate Hh signalling in response to pur, but that treatment was not sufficient to restore muscle regeneration.

**Figure 6:**
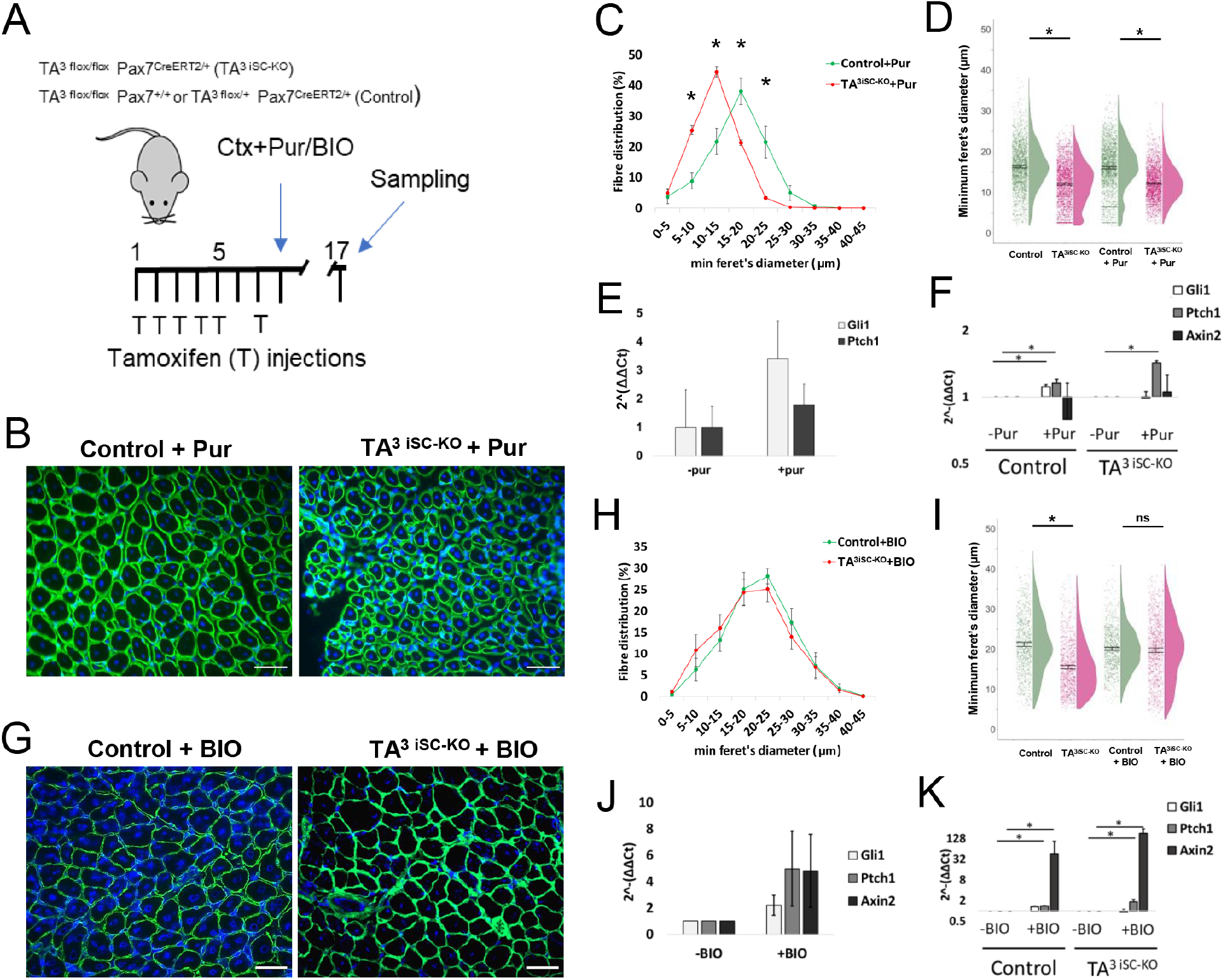
GSK3ß inhibition restores muscle regeneration. (A) Schematic of the experimental approach. (B, G) Transverse sections ten days post-injury stained for nidogen (green) and DAPI (blue). (B) Sections are shown for control and TA^3iSC-KO^ muscles, co-injected with ctx and purmorphamine (pur), or (G) co-injected with ctx and BIO (scale bar = 50 μm). (C, D, H, I) Histograms and raincloud plots show the range of myofiber diameters in control (green) or TA^3iSC-KO^ (red) mice, injected with ctx and pur (C, D) or with ctx and BIO (H, I). (E) qRT-PCR shows expression of Gli1 and Ptch1 in whole muscle, with or without pur. (F) qRT-PCR shows fold change of expression in FACS isolated MuSC progeny for Gli1, Ptch1 and Axin2, with or without pur. (J) qRT-PCR shows expression of Gli1, Ptch1 and Axin2 in whole muscle, with or without BIO. (K) qRT-PCR shows fold change of expression in FACS isolated MuSC progeny for Gli1, Ptch1 and Axin2, with or without BIO. * indicates p≤0.05 calculated by unpaired, two-tailed t-test. n?3 mice.

GSK3ßkinase is a major negative regulator of ß-catenin dependent (canonical) Wnt signalling. It is highly expressed in muscle and inhibition of GSK3ßenhances self-renewal of embryonic stem cells ^30^. Given that expression of Wnt ligands is reduced in TA^3iSC-KO^ MuSCs we examined the effect of GSK3ßinhibition. Muscle regeneration was restored after co-injection of ctx with BIO (6- bromoindirubin-3’-oxime) and myofiber diameters were similar in TA^3iSC-KO^ and control mice, indicating that BIO could ameliorate the regeneration defect in absence of primary cilia (Fig. 6G-I). Since BIOmediated GSK3ßinhibition activates ß-catenin dependent Wnt signalling and Hh signalling, we assessed expression of Axin2, a Wnt target gene, alongside Gli1 and Ptch1. In whole muscle biopsies, expression of all three marker genes was increased by BIO (Fig. 6J). In FACS isolated MuSCs, application of BIO induced expression of Axin2 and to a lesser extent the expression of Gli1 and Ptch1 (Fig. 6K). This suggests that MuSCs activate predominantly ß-catenin dependent Wnt signalling in response to BIO and identifies canonical Wnt activation as a treatment to overcome the loss of cilia.

## Discussion

Quiescent muscle stem cells have a primary cilium, which is dynamically regulated during the myogenic cycle and is reassembled in self-renewing stem cells ^17,18,31,32^. Furthermore, myofibers isolated from aged muscles have fewer MuSCs with a primary cilium, indicating that cilia might be lost with age or become less stable ^19^. Ciliopathies, a group of genetic disorders resulting from perturbation of ciliary proteins in human, affect several different tissues including muscle. For example, Joubert Syndrome, which can result from mutations in the Talpid3 gene (KIAA0586) ^10^, is associated with hypotonia ^33, 34^. Here, we examine the role of TA^3^ in MuSCs and use inducible deletion to show that primary cilia are essential for efficient regeneration after muscle injury *in vivo*.Using single cell sequencing of MuSCs, we found that both Hh and Wnt signalling were deregulated in absence of TA^3^. Importantly, we discovered that pharmacological activation of Wnt signalling, but not Hh signalling, can restore muscle repair in the absence of a primary cilium. These findings provide insights into potential avenues for treatments in muscle disease or ageing.

It has previously been proposed that the primary cilium may provide a potential target for regenerative therapies ^19, 21, 35^ and recent work has implicated the cilium in muscle regeneration. Specifically, deletion of the intra-flagellar transport protein, IFT88, in MuSCs led to a shift toward myofibers with reduced cross-sectional area post-injury ^19^. This agrees with our finding that absence of primary cilia impairs muscle repair and that, following injury, regenerating TA^3iSC-KO^ myofibers had smaller diameters compared to controls. In addition, we found that repeat injury enhanced the regeneration defect and led to a more severe phenotype (Fig. 4). This suggests, MuSCs are depleted and their expansion and self-renewal could be affected in absence of TA^3^. It is striking that MuSCs lacking GLI3, which acts predominantly as a repressor, display rapid cell-cycle entry, increased proliferation and expansion of the stem cell pool ^21^, whereas loss of cilia inhibits proliferation and MuSC expansion. This was observed after deletion of IFT88 in MuSCs ^19^ and in this report where following TA^3^ deletion the number of MuSC progeny was reduced on *ex vivo* cultured myofibers (Fig. 2). This suggests the cilium is necessary to maintain an appropriate balance of GLI activator and repressor proteins ^20^.

Molecular profiling of individual MuSCs showed that expression of cilia related genes was significantly reduced in cells lacking TA^3^ (Fig. 5E). UMAP clustering identified a population of MuSCs positive for Pax7. This population could be sub-divided into two clusters, one of which was characterised by the expression of genes related to cell division, such as Cdk1 (Fig. 5D) and comprised mitotically active, proliferating cells. The second cluster of MuSCs positive for Pax7 was enriched for genes associated with ECM related processes. We propose that this cluster comprises MuSCs that are close to quiescence, similar to those recently described after single cell sequencing of whole hindlimb muscle ^36^. This cluster may also include self-renewed MuSCs that have entered quiescence, as some cells express low transcript levels for myogenic regulatory factors, such as MyoD (Fig. 5E) or Myf5.

During embryogenesis, Sonic hedgehog signalling regulates basement membrane components, such as laminin-111 ^(37–39^. This is crucial for myotome formation and basement membranes are disrupted in the TA^3^ mutant ^10^. It is therefore interesting to note that TA^3^ deletion led to differences in expression of ECM related genes, in quiescent MuSC isolated immediately following the deletion of TA^3^. This included tissue inhibitor or matrix metalloprotease 1 (TIMP1), matrillin 2 (Matn2), laminin subunit beta 1 (LAMB1), integrin alpha 8 (Itga8) and ADAM Metallopeptidase Domain 12 (ADAM12), which were all upregulated in absence of TA^3^. However overall, there were few significant differences in MuSC transcriptomes at this time point (Supplemental Fig. 4). This is not unexpected as these cells are transcriptionally not very active. The differences we do see may be due to partial activation induced by the isolation process ^(40–43^.

As primary cilia are implicated in both Hh and Wnt signalling, we examined the expression of components of these pathways in control and TA^3iSC-KO^ MuSCs. Differential analysis showed that both pathways were deregulated in absence of TA^3^ (Fig. 5F). During regeneration, canonical Wnt signalling is activated and has been implicated in regulating myogenic lineage progression ^44^. However, ß-catenin dependent Wnt signalling is only transiently active in expanding myoblasts, and its restriction is crucial for successful muscle regeneration ^45^. Wnt signalling is also important in embryonic myogenesis ^(46–48^ as well as during terminal differentiation, where it activates follistatin and myoblast fusion ^49^. Furthermore, Sonic hedgehog synergises with Wnt to activate myogenesis in paraxial mesoderm progenitors in both mouse and chick ^(50–52^. Thus, the role of these signalling pathways in myogenesis is complex.

It was particularly striking that Hedgehog interacting protein (Hhip), Patched2 (Ptch2) and several Wnt ligands, Wnt10a, Wnt5a, Wnt9a, Wnt2 and Wnt7b, were significantly reduced in TA^3iSC-KO^MuSCs (Fig. 5F). This observation prompted us to attempt pharmacological rescue of the regeneration defect. Co-injection of ctx and purmorphamine (pur), a smoothened agonist ^25^, activated Hh target genes in MuSCs, however did not restore muscle repair (Fig. 6B-F). This agrees with a previous report, where injection of SAG1.3, another Hh pathway agonist, had no effect in mice with IFT88^-/-^ MuSCs, either on muscle strength or MuSC proliferation ^19^. Thus, the Hh mediated effects on MuSC during the regenerative process appear to be completely cilia dependent. In contrast, co-injection of ctx with an inhibitor of GSK3ßkinase (BIO) activated expression of Axin2, a canonical Wnt target gene, in MuSCs and restored muscle repair independenly of primary cilia (Fig. 6G-K). GSK3ßkinase is highly expressed in muscle; it regulates many signalling pathways including Shh, Wnt and Notch. It has been shown that BIO mediated suppression of GSK-3 activity promotes Wnt/beta-catenin signalling to preserve the pluripotency of human and mouse embryonic stem cells ^30^. GLI2, which is part of the Hh pathway, can also be phosphorylated by GSK-3 ^53, 54^, however, BIO did not significantly activate the expression of Hh target genes in MuSCs. GSK-3β can also regulate the stability of surface Notch receptors and there is extensive cross-talk between Wnt and Notch signalling ^55, 56^. However, our transcriptomics analysis did not identify differentially expressed Notch pathway components.

In summary, we show that BIO restores muscle regeneration and that this works predominantly through activation of Wnt signalling, which can overcome the lack of TA^3^ and primary cilia. Thus, our study identifies a novel therapeutic target in MuSCs that could be of interest in muscle degenerative disease or in ageing.

## Experimental Methods

### Animal models

All mice were housed in the Disease Modelling Unit (DMU) of the University of East Anglia. Experimental procedures were performed in accordance with the Animal (Scientific Procedures) Act 1986 (ASPA) under the UK Home Office Project Licence PP3253888. All lines were maintained on a C57BL/6J background: Pax7^CreERT2/+ 23^, Talpid3^flox/flox 12^. B6.Cg-Tg(Pax7-ZsGreen)^1Kyba/J^ (kindly provided by M. Kyba, University of Minnesota, USA ^24^; and Rosa26-td-Tomato (Ai14, The Jackson Laboratory). Mice carrying the TA^3^ floxed allele were crossed with mice expressing the tamoxifen inducible cre-recombinase, CreERT2, under the control of the endogenous Pax7 promoter (Pax7^CreERT2^), generating control mice and TA^3iSC-KO^ in the same litter. Experimental mice were 8-12 weeks old. For genotyping DNA was extracted from ear notches and polymerase chain reaction (PCR) was performed as described ^12,23^.

### IP injections

Tamoxifen was injected for five consecutive days, followed by a rest day and another tamoxifen injection on day seven. During the regeneration phase tamoxifen was administered every other day.

### Muscle injury model

Mice were anaesthetized with isofluorane and muscle injury was induced with 50μl of a 10μM cardiotoxin (ctx) (Latoxan) solution in the right *tibialis anterior* muscle. The un-injured left leg served as control. Muscles were isolated at the indicated time points after injury. For double injury experiments, a second injury was induced after 25 days of recovery from the first injury. Muscle was then analysed 10 days post repeat-injury. For the rescue experiments purmorphamine (2 μM) or BIO (10 μM) were injected together with ctx.

### Histology and Immunofluorescence

To confirm the degree of injury and regeneration, at the given time points mice were sacrificed and skeletal muscles were harvested, fixed in 4% PFA, embedded in OCT and transverse cryosectioned (10μm), followed by Hematoxylin-Eosin. For immune staining sections were blocked with 5% Normal Goat Serum in TBS-T, followed by incubation with Nidogen1 (Willem et al., 2002), after washing Alexa Fluor 488 secondary antibody and DAPI (0.5μg/ml) were applied. Slides were mounted with gelvatol for imaging.

### Muscle fiber isolation

Following administration of tamoxifen *extensor digitorum longus* (EDL) muscles were isolated from 8-12 week-old mice and incubated in Collagenase I (2mg/ml, Worthington,) for 90 minutes at 37°C, after which muscles were transferred to BSA (Sigma) coated dishes containing DMEM (Gibco) and PenStrep (Gibco). EDL muscles were subjected to repeated trituration through descending diameter glass Pasteur pipettes until single myofibers separated from the isolated muscle. Single myofibers were selected and cultured in 10% Horse Serum (HS), 1% Chick Embryo Extract (CEE) and PenStrep in DMEM.

### Immunofluorescence

Transverse sections were blocked with 5% Normal Goat Serum (NGS) in Tris-buffered saline containing 0.1% Tween 20 (TBS-T) for 60 min at 37°C, followed by a 60 minute incubation with Nidogen1 as primary antibody (Willem et al., 2002). After three 5 minute washes in TBST-T, sections were incubated with Alexa Fluor 488 secondary antibodies and DAPI (0.5μg/ml) in 2% NGS/TBS-T for 60 min at room temperature, washed three times in TBS-T and mounted with gelvatol.

At indicated time points, muscle fibers were fixed for 10’ with prewarmed 2% Paraformaldehyde (PFA), washed with PBS and permeabilised by adding 0.5% TritonX100 and blocked for 2hrs with 10% NGS in TBS-T at 37°C. Primary antibodies against Pax7 (DSHB), Myf5, MyoD (Santa Cruz) and Myogenin (DSHB) were used at a 1:50 dilution in 5% NGS/TBS-T, incubated at 4°C overnight. Samples were extensively washed with TBS-T before secondary antibodies (Jacksonimmuno) were applied together with DAPI for 2 hours in the dark. Fibers were washed in TBS-T and TBS, and 10-20 fibers were mounted in gelvatol on a glass slide for imaging.

For immunostaining of primary cilia fixed fibers were permeabilized with 0.5% TritonX100 and 0.5% Nonidet P40 and 2% MeOH. Primary antibodies used were directed against Arl13B (Proteintech, 1:300) and Pax7 (DSHB; 1:7.5). Secondary antibodies used were goat anti-mouse IgG-Fab2 488 and goat anti-rabbit Alexa fluor 546 (1:375) (Jacksonimmuno).

### Microscopy and image analysis

Fluorescence images of muscle sections and muscle fibers were taken using a Zeiss AxioPlan 2ie microscope. Images of muscle sections were processed in Fiji (ImageJ) after being pixel corrected in Ilastik and the minimum Feret diameter was measured.

### Fluorescence-activated cell sorting

Quiescent MuSCs (Pax7-ZsGreen+) were obtained from uninjured hindlimb muscles. For scRNAseq cells were isolated from *ex vivo* cultured myofibers (72 hrs) and Td-Tomato positive MuSCs were sorted using the BD FACSMelody cell sorter (BD Biosciences, San Jose, California) directly into lysis buffer in 96-well plates. From three independent experiments we recovered two plates each from control and TA^3iSC-KO^ myofibres, generating 6 plates per genotype.

### Single-cell RNA-seq

Amplified cDNA was generated from sorted cells using the SmartSeq2 protocol (Picelli et al, 2014). Sequencing libraries were generated using an automated, reduced-volume version of the Nextera XT protocol (Illumina) using the SPT Labtech Mosquito LV. Libraries were pooled (384-plex) and sequenced on a NovaSeq 6000 (Illumina, San Diego) in paired end, dual index mode. Raw Illumina sequencing data were analysed to obtain a single-cell expression matrix object. Subsequent analysis was performed in R using Seurat version 3 ^28^. Cells showing gene counts lower than 1,000 and a mitochondrial gene expression percentage higher than 5% were excluded from further analysis. Within Seurat, data were normalised using NormalizeData (normalization.method =“LogNormalize”, scale.factor = 10000) and data from multiple samples were merged using the FindIntegrationAnchors and IntegrateData commands.

### Statistical analysis

A minimum of three independent experiments (or animals) was used for all asssys. Statistical analysis was carried out using Excel software. Multiple data points were collected per mouse to generate the population mean and represent it as raincloud plots using ggplot2 in R-studio. These values were used to compare between control and mutant mice. Unpaired, two tailed students T-tests pairwise were used to calculate statistical significances, shown as P values. Histograms were generated using Microsoft Excel and comparison was done at each fiber range.

### RNA extraction and real time qPCR

RNA from the Td-Tomato positive cells was extracted using trizol and qPCR was performed in triplicates by using the RNA to CT-1-step Sybr green kit (Life Technologies Ltd). Primers to detect transcripts of Hh and Wnt targets are in Supplementary Table 1.

### Data availability

The raw sequencing data can be accessed on the NCBI-SRA archive, accession number xxx (to be confirmed).

## ACKNOWLEDGEMENTS

We thank Paul Thomas in the Henry Wellcome Cell Imaging Laboratory (UEA) for support with microscopy, Anita Scoones (Earlham Institute) for discussions of scRNA analysis, Peter Zammit (KCL) for sharing advice on myofiber isolation, and Cheryll Tickle for sharing the TA^3^ floxed mice. VMH, LF, SS and TP were supported by MRC project grant, MR/R000549/1, to AM and UM. DB was supported by a BBSRC NRPDTP studentship. GFM was supported by BHF project grant, PG/19/76/34696. The authors acknowledge funding from the Biotechnology and Biological Sciences Research Council (BBSRC), part of UK Research and Innovation, Core Capability Grant BB/CCG1720/1. Part of this work was delivered via the BBSRC National Capability in Genomics and Single Cell Analysis (BBS/E/T/000PR9816) at Earlham Institute.

## AUTHOR CONTRIBUTIONS

AM and UM conceived the project, designed experiments, discussed results and provided financial support. VMH, DB, SS and TP conducted experiments and analysed results. LF processed and analysed bulk RNAseq data. GFM analysed scRNA-seq data. IMcC, LM and EP helped with FACS isolation. IMcC and LM made libraries for scRNA-sequencing with assistance from CU. AM wrote the draft manuscript with input from co-authors. All authors discussed and edited the final manuscript.

## Supplementary Figures and Tables

**Supplementary Table 1.**
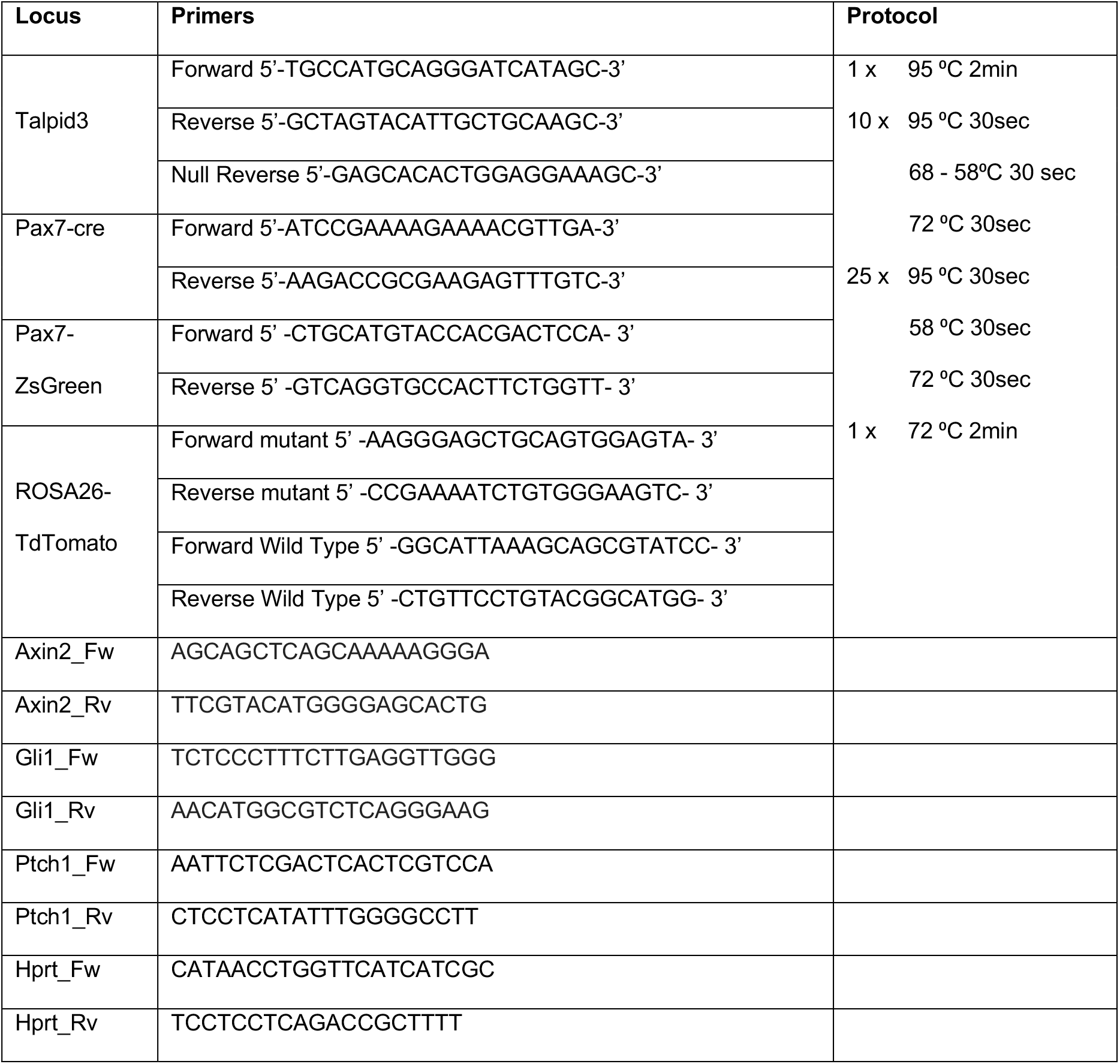
Primers and protocols for mouse Genotyping.

**Supplementary Figure 1:**
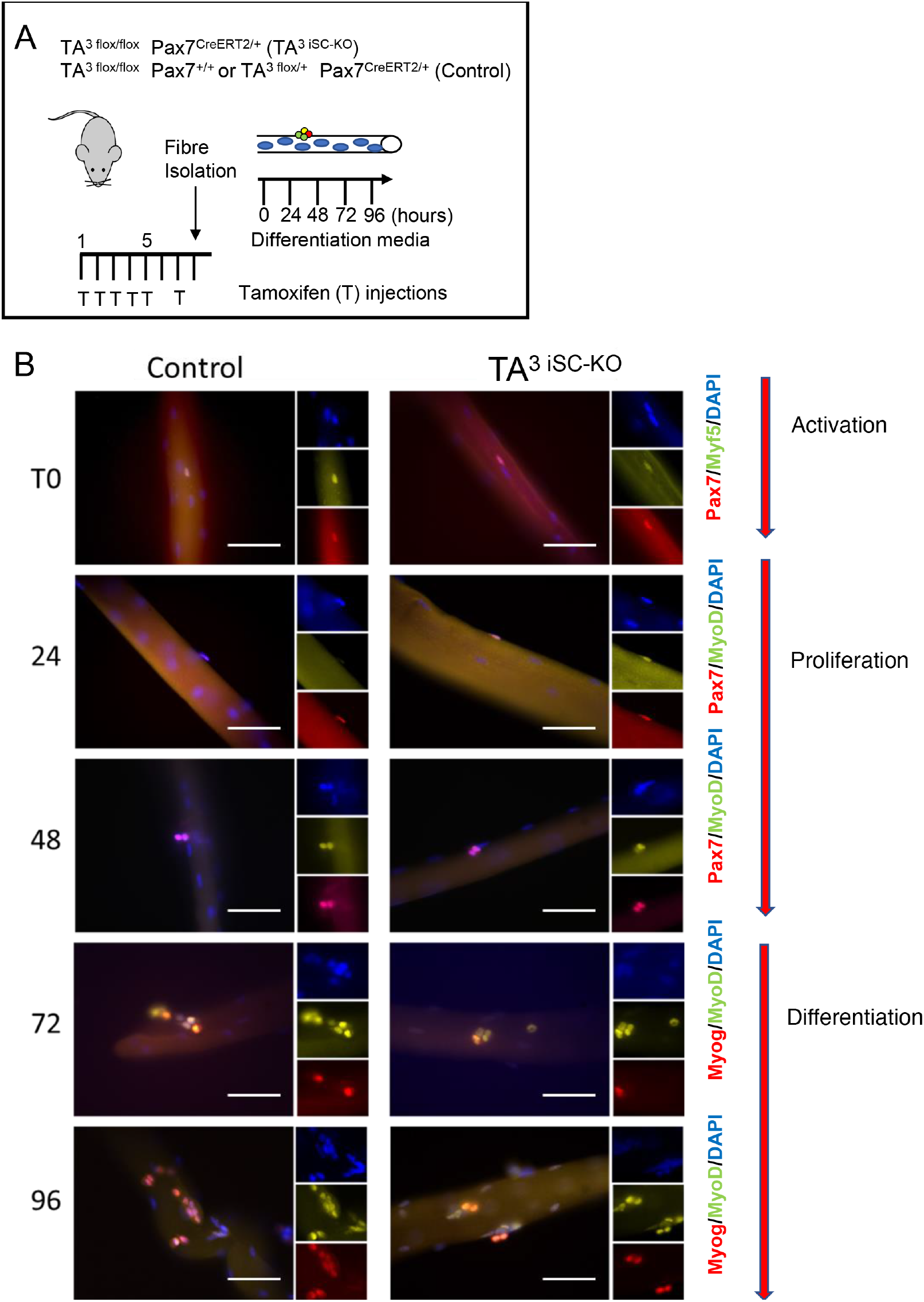
Immunostaining of MuSC on *ex vivo* cultured myofibers with different marker genes. (A) Schematic representation of the experimental design. (B) Representative images of EDL muscle fibers from control and TA^3iSC-KO^ mice expressing markers characteristic for each stage of the regenerative cycle (size bars = 50 μm).

**Supplementary Figure 2:**
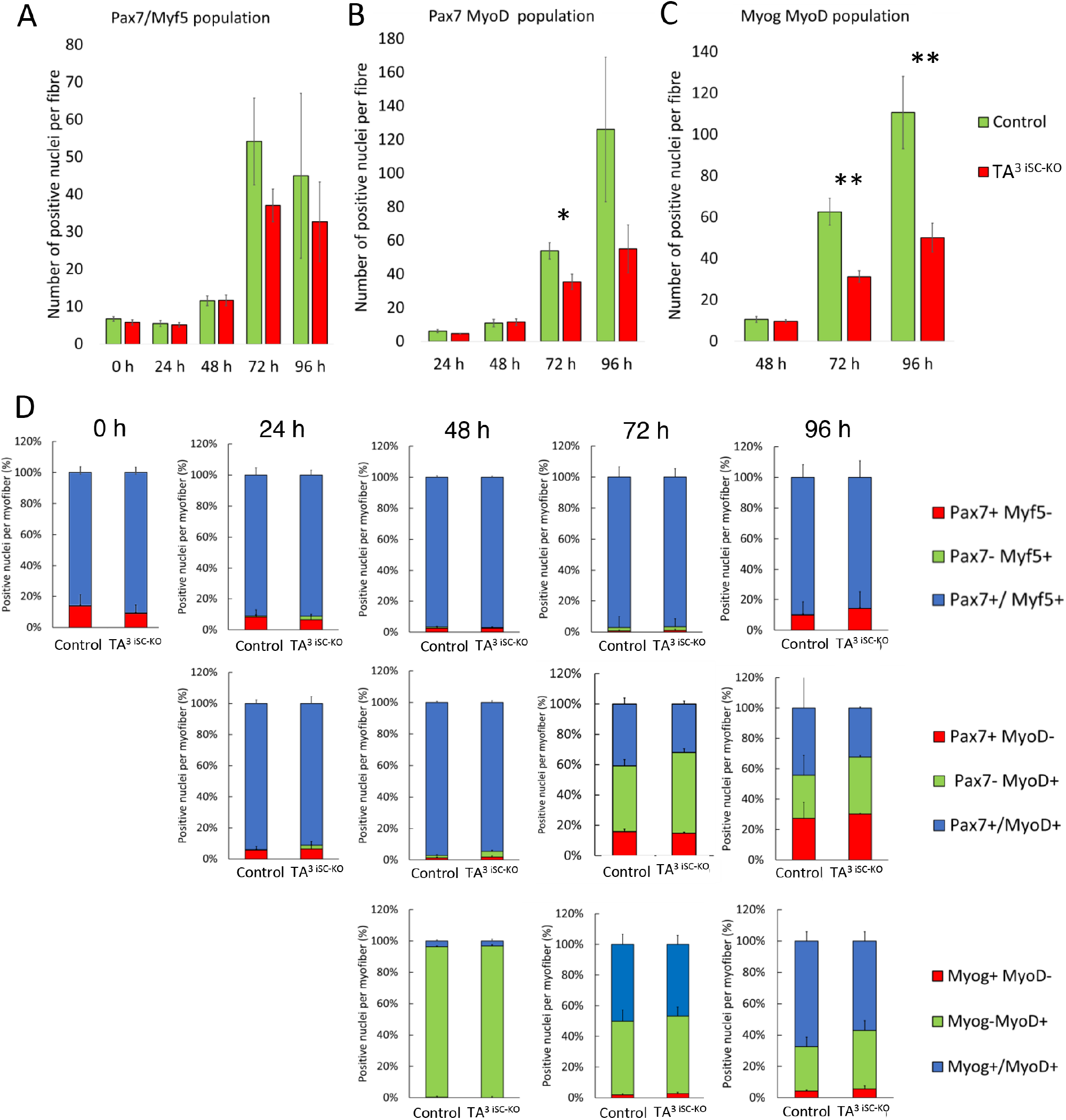
Quantification of MuSC immunostaining on *ex vivo* cultured myofibers. (A) Total number of cells that express Pax7 and/or Myf5, (B) total number of cells that express Pax7 and/or MyoD, (C) total number of cells that express MyoG and/or MyoD at the time points indicated. Controls (green) and TA^3iSC-KO^ (red), n=9. *p<0.05, **p<0.01 (D) Percentage of cells that are positive for one or both of the markers indicated at the different time points in myofibers from controls and TA^3iSC-KO^ mice, n=9, *p<0.05, **p<0.01

**Supplementary Figure 3:**
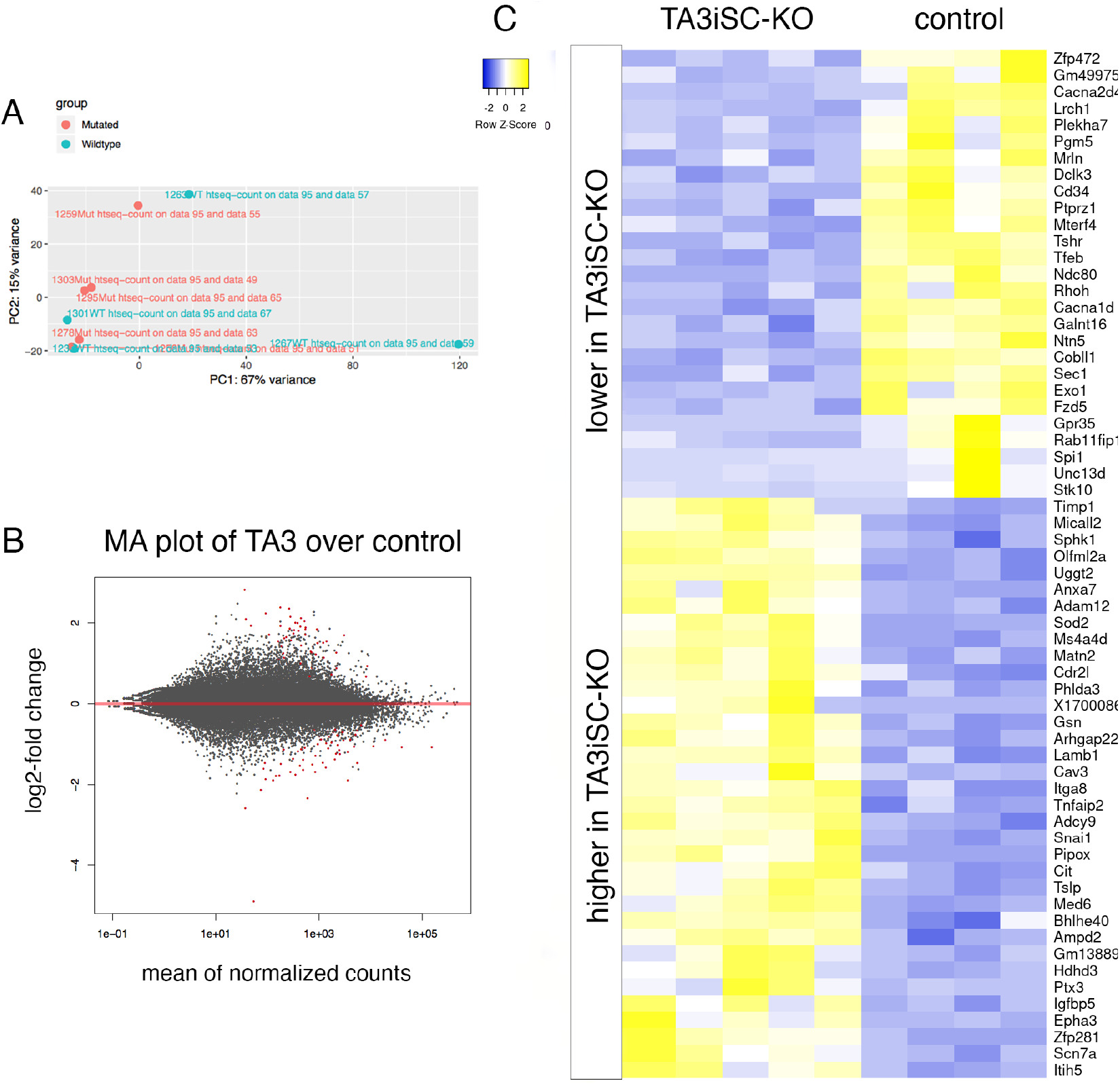
Pax7-ZsGreen positive MuSCs isolated by FACS from control and TA^3iSC-KO^ mice have similar molecular profiles immediately following deletion. (A) PCA plots of RNAseq samples from control (blue) and TA^3iSC-KO^ MuSCs (red). (B) MA plots showing log2-fold change of differentially expressed (DE) genes. (C) Heatmap of DE genes, p-value <0.05.

**Supplementary Figure 4.**
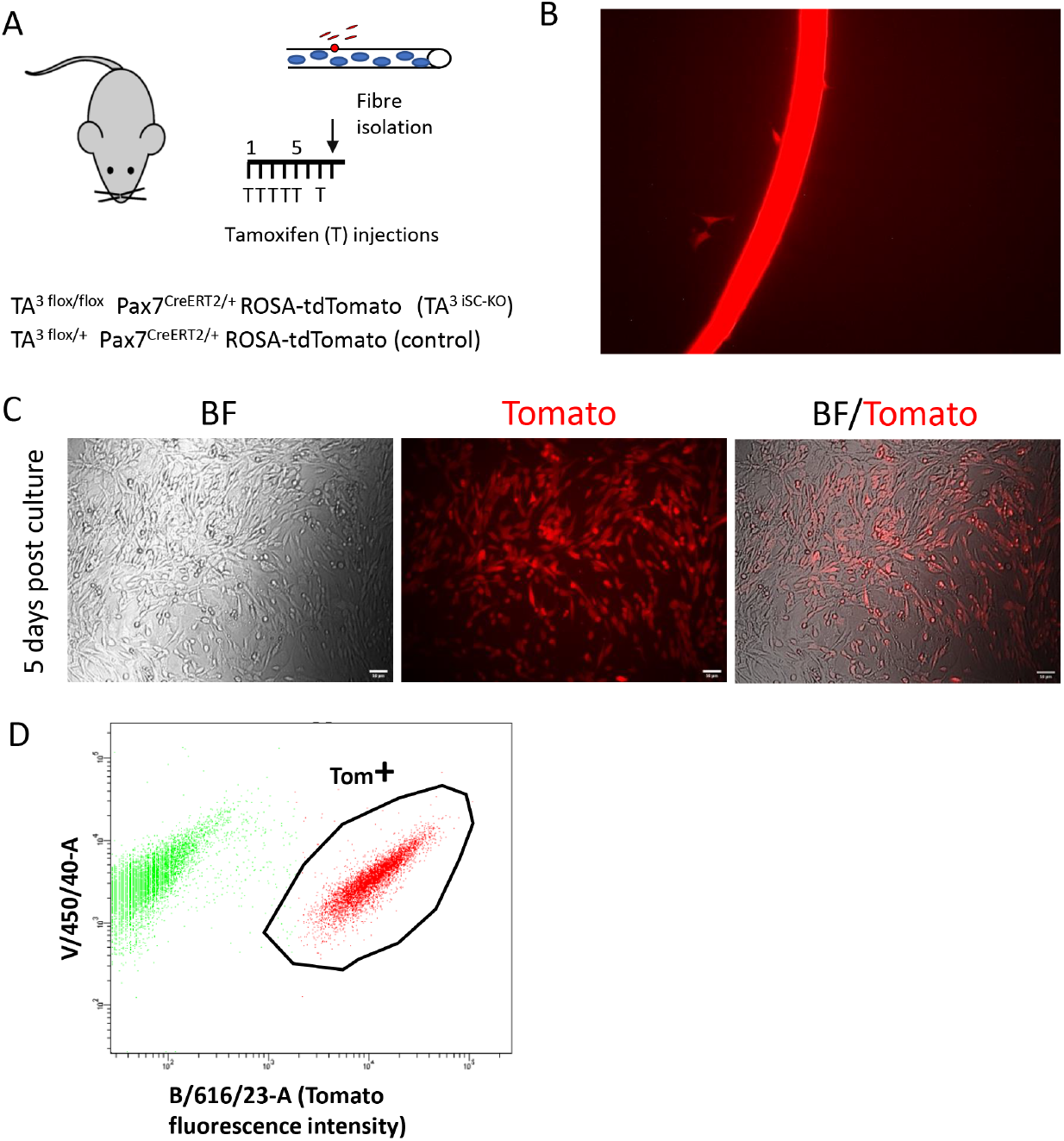
FACS isolation of MuSCs. (A) Schematic representation of the experimental design. (B) MuSCs from cultured fibers grow out on matrigel coated plates. (C) MuSC progeny expanding prior to drug treatments and FACS, BF, bright field; Tomato, Td-Tomato fluorescence. (D) Scatter plot showing Td-Tomato positive cells separating from the rest of the population.

